# Realistic Anatomically Detailed Open-Source Spinal Cord Stimulation (RADO-SCS) Model

**DOI:** 10.1101/857946

**Authors:** Niranjan Khadka, Xijie Liu, Hans Zander, Jaiti Swami, Evan Rogers, Scott F. Lempka, Marom Bikson

## Abstract

**Objective:** Computational current flow models of spinal cord stimulation (SCS) are widely used in device development, clinical trial design, and patient programming. Proprietary models of varied sophistication have been developed. An open-source model with state-of-the-art precision would serve as a standard for SCS simulation.

**Approach:** We developed a sophisticated SCS modeling platform, named Realistic Anatomically Detailed Open-Source Spinal Cord Stimulation (RADO-SCS) model. This platform consists of realistic and detailed spinal cord and ancillary tissues anatomy derived based on prior imaging and cadaveric studies. Represented tissues within the T9-T11 spine levels include vertebrae, intravertebral discs, epidural space, dura, CSF, white-matter, gray-matter, dorsal and ventral roots and rootlets, dorsal root ganglion, sympathetic chain, thoracic aorta, epidural space vasculature, white-matter vasculature, and thorax. As an exemplary, a bipolar SCS montage was simulated to illustrate the model workflow from the electric field calculated from a finite element model (FEM) to activation thresholds predicted for individual axons populating the spinal cord.

**Main Results:** Compared to prior models, RADO-SCS meets or exceeds detail for every tissue compartment. The resulting electric fields in white and gray-matter, and axon model activation thresholds are broadly consistent with prior stimulations.

**Significance:** The RADO-SCS can be used to simulate any SCS approach with both unprecedented resolution (precision) and transparency (reproducibility). Freely available online, the RADO-SCS will be updated continuously with version control.

## Introduction

### Broad impact of an open-source high-resolution computational SCS model

Computational models predict current flow patterns and neuronal activation during neuromodulation techniques, such as spinal cord stimulation (SCS) ^1–3^. These models are key tools in designing, optimizing, and understanding SCS as they relate the *controllable* stimulation dose (i.e. electrode placement and waveform ^4^) with the *intended* resulting activation of the spinal cord and nerves ^5,6^. Computational SCS models thus broadly inform modern clinical SCS practices, ongoing research into mechanisms of actions, and design of new interventions ^2,7–9 10–14^.

Models of SCS have been continuously refined and applied from the early 80’s through recent efforts ^1,2,5,10,13–31^ (see Table 1). Development of models of increasing complexity offered mirrored general enhancements in numerical modeling techniques (finite element analysis), with proprietary efforts by numerous groups, each subject to multiple version iterations. Without open-source model-geometry and a standard modeling pipeline, exact replication is difficult. Indeed, the more advanced (detailed) a model, the more intractable the model is to reproduce without source code. Moreover, even recent models can lack details of major anatomical structures of the spine.

**Table 1:**
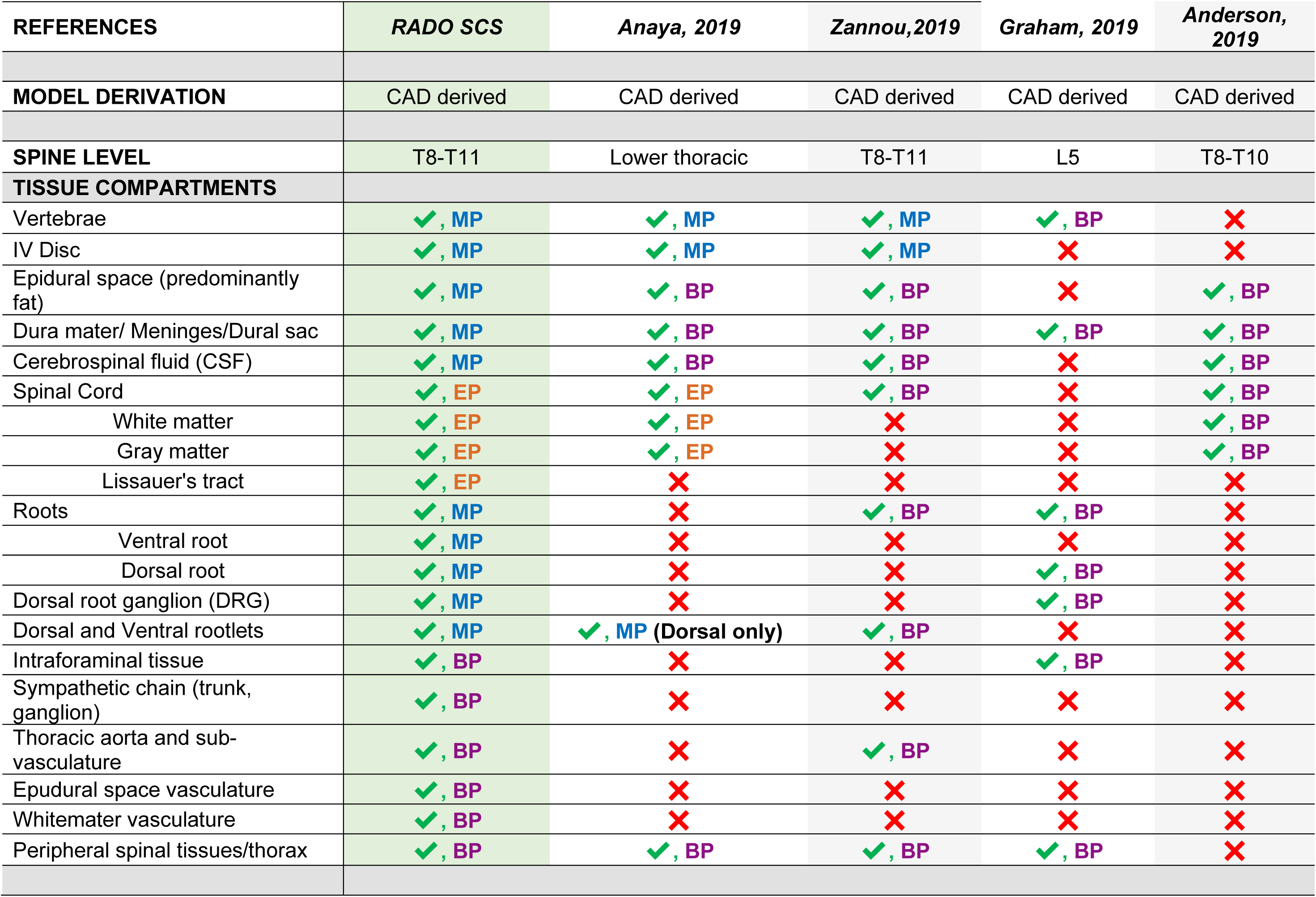
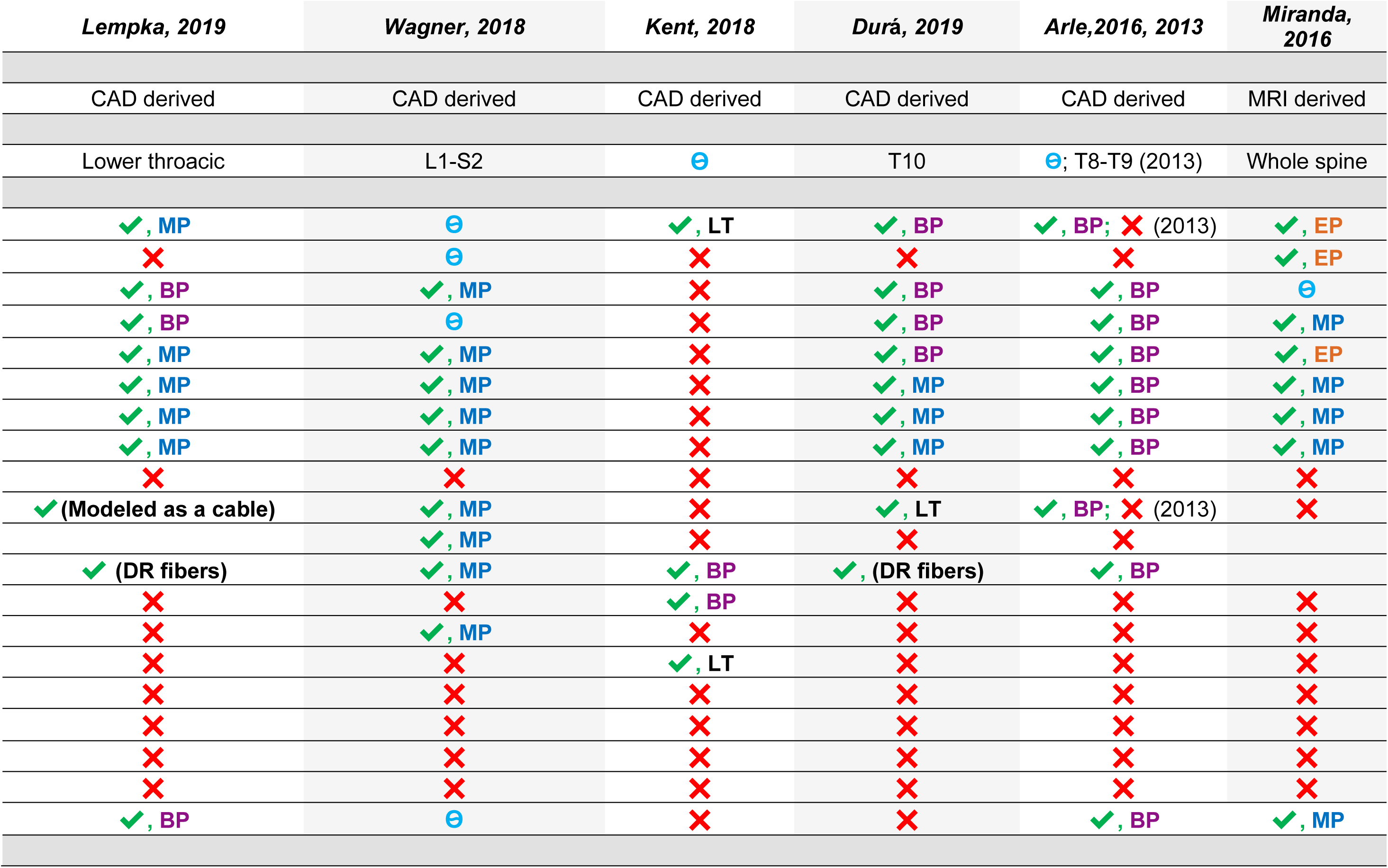
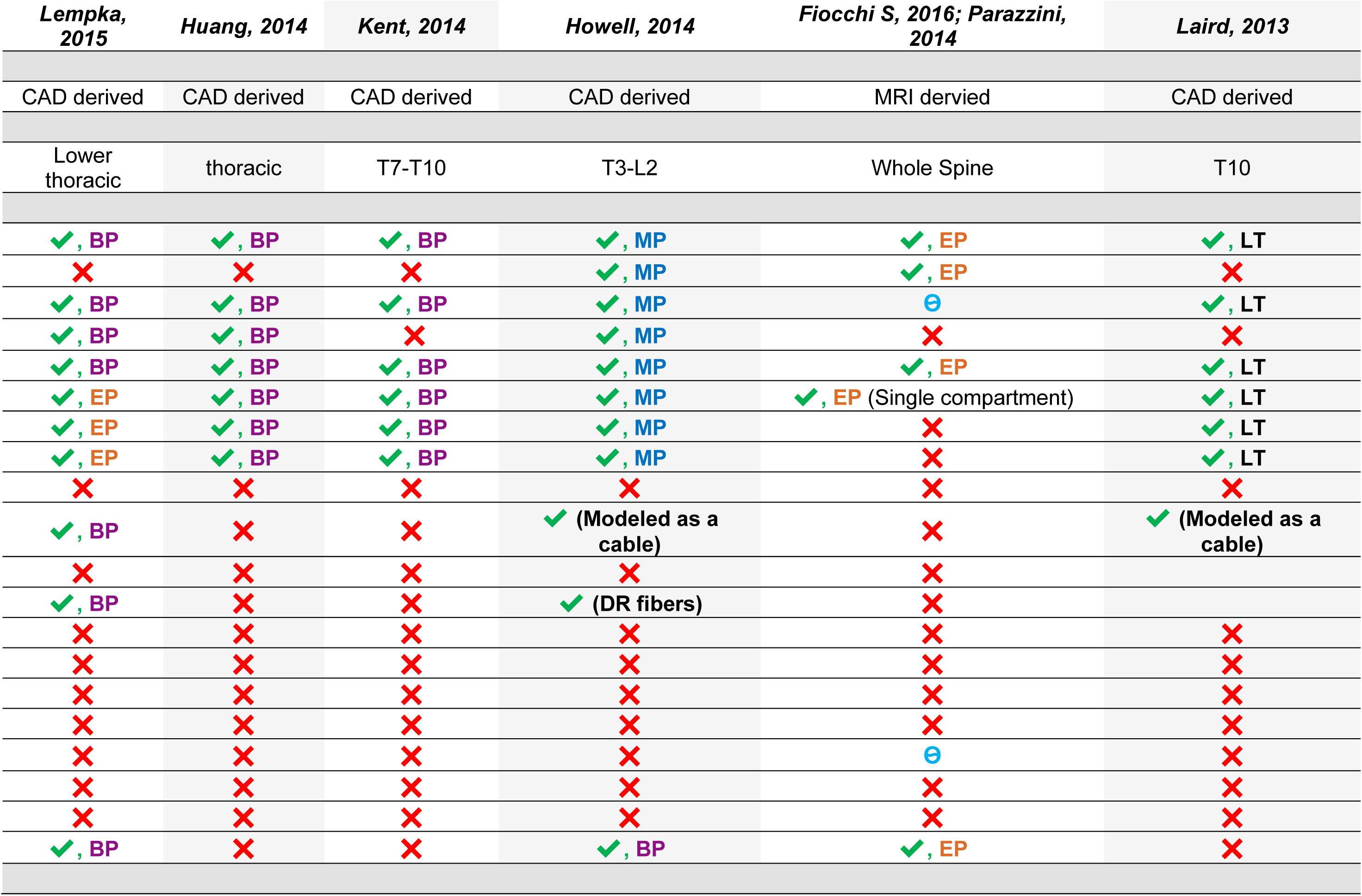
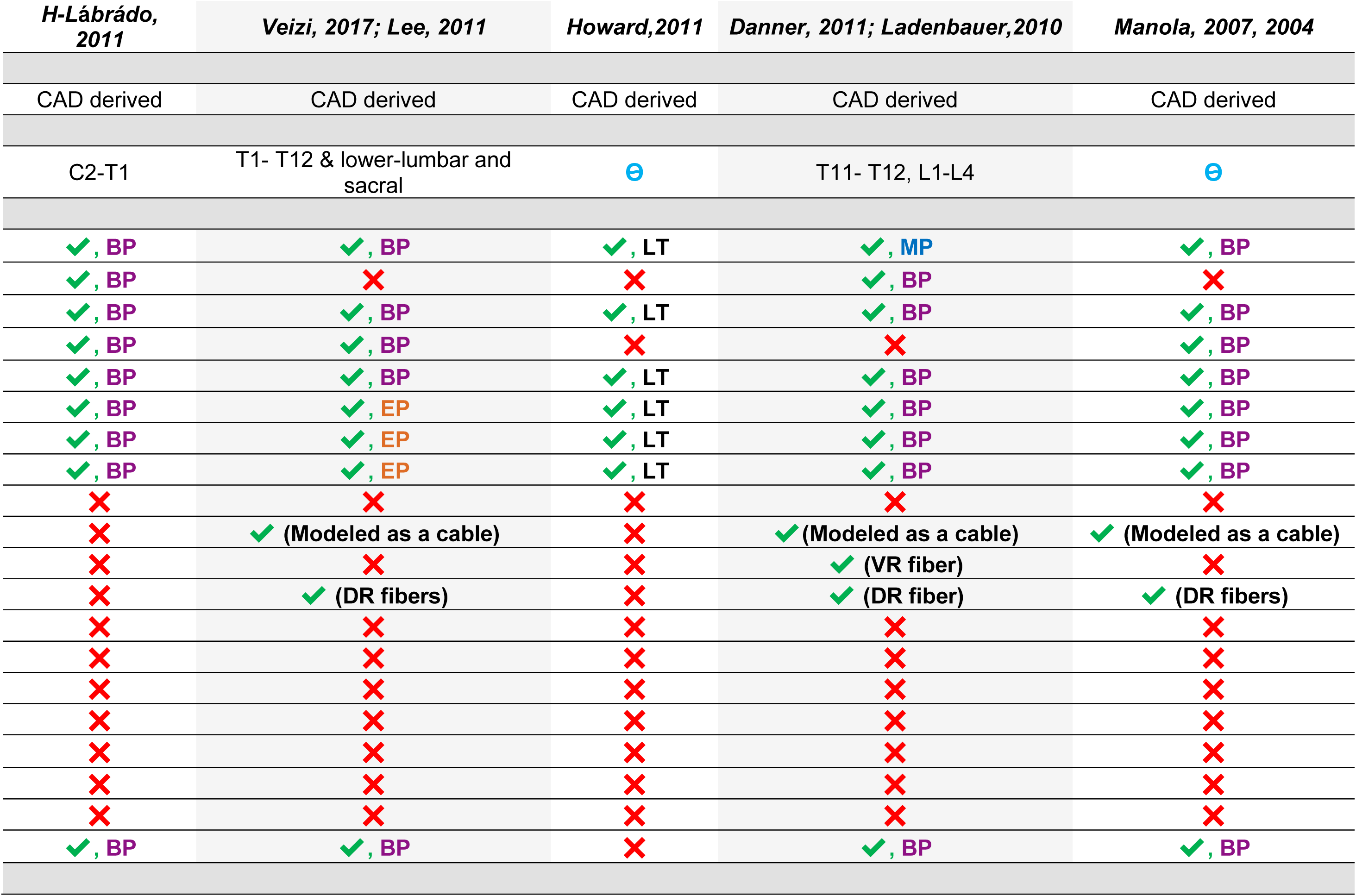
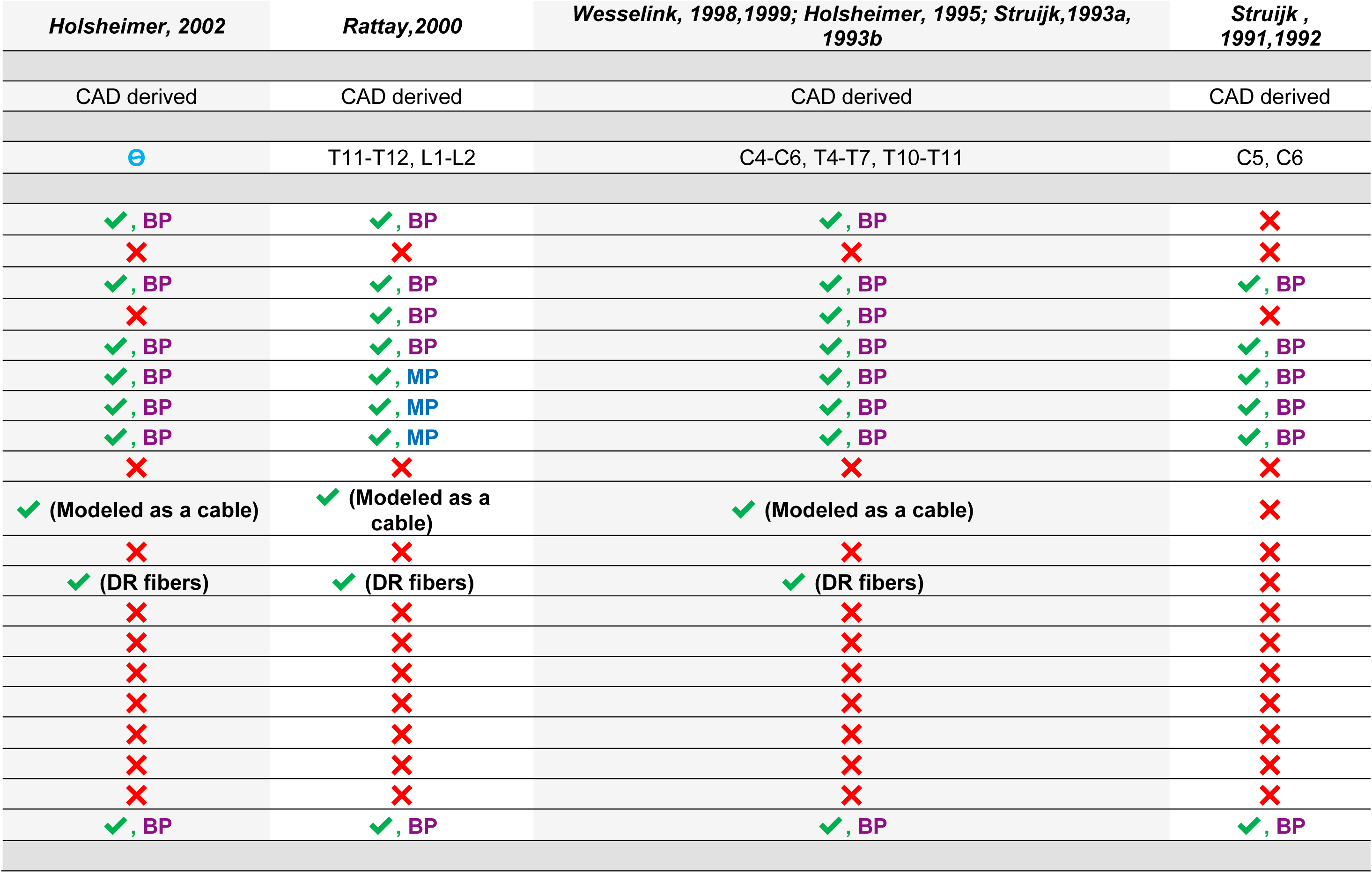
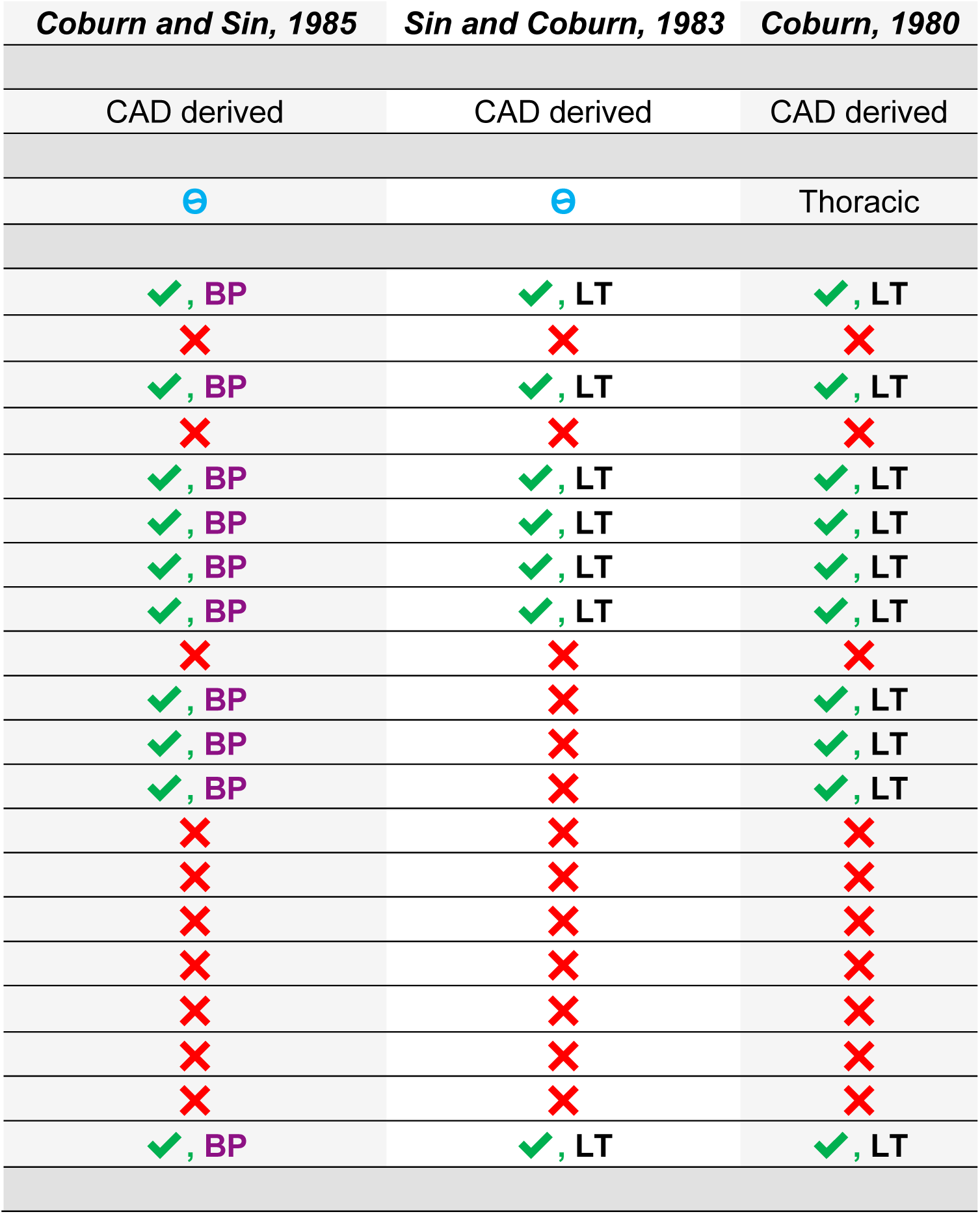
Comparison of the RADO SCS model with other existing SCS models based on model derivation (CAD vs. MRI), constructed tissue compartments (unclear, considered, and not considered/absent), and precision in anatomical details (limited, basic, moderate, and enhanced).

Here, we develop the first Realistic Anatomically Detailed Open-source Spinal Cord Stimulation (RADO-SCS) model: all assembled CAD files (STL) of spinal tissues, along with available devices renders, meshes, FEM results, and activation simulations are available for free download under an open-source license. RADO-SCS supports stimulation of any SCS dose (technology) and will be subject to ongoing updates with version control. The more precise and complex a computational model, the more critical it is to share code for reproducibility and to prevent the need to redo the resource-intensive creation effort. Use of RADO-SCS thus provided users with 1) a transparent and reproducible platform to base any claims; 2) evolving state-of-the-art precision to best model quality; and 3) cost and time savings. RADO-SCS is a unique tool for supporting computer-driven device design, dose optimization, and efficient clinical trial design.

### Anatomical details of prior SCS models

Finite element analysis has been widely implemented in two-dimensional (2D) and three-dimensional (3D) spinal cord current flow models. Although multitude computational SCS models of rodent and non-human primates had been developed and implemented for motor control following spinal cord injury ^7,32,33^, here we specifically focused on human SCS modelling studies (minimally invasive or non-invasive) for pain management. We categorized prior SCS models based on tissue compartments (considered vs. not considered/absent) and anatomical precision (limited, basic precision, moderate precision, and enhanced precision) (see **Table 1**). Limited refers to SCS models with minimal anatomical precision in constructed tissue compartments. Basic precision SCS models have regular shapes (e.g., cylindrical, triangular, rectangular prism, wedge, bricks, or cube) as tissue compartments, no flexion in geometry, and uniform dimension across spine levels. Moderate precision SCS models include tissue compartments with minimal resolution or have regular geometric shapes with some flexion and uniform dimension across spine level. Enhanced precision refers to SCS models with tissue compartments with realistic geometry, additional anatomical details (flexion, bifurcation, union), high resolution, and spine level specific dimensions.

In the early 80’s, Coburn developed the first 2D FEM model representing non-homogeneous human spinal cord tissues for noninvasive and invasive SCS. The model comprised major spinal tissue domains, such as thoracic vertebrae, epidural fat, CSF, spinal roots, white-matter, and gray-matter with limited precision ^17^. In 1983, Sin and Coburn developed a simplified version of the Coburn1980 2D SCS model, excluding spinal roots (limited precision) ^34^. Coburn and colleagues in 1985, and Struijk and colleagues in 1991 and 1992 developed 3D FEM models of SCS including major tissue of the spinal canal with basic precision and coarse meshing. In both models, dura was absent, with only Coburn’s model including a basic spinal root ^18,29,35,36^. Struijk and colleagues in 1993a and 1993b, Holsheimer and colleagues in 1995, and Wesselink and colleagues in 1998 and 1999 developed a 3D SCS model including the mid cervical (C4-C6), mid thoracic (T4-T7), and low thoracic (T10-T11) vertebral spine level with major spinal canal tissue compartments, including dura mater and surrounding tissue layers (thorax)-all tissue compartments with basic precision ^28,14,31,37,38,12^. Wesselink et al. included white matter anisotropy and encapsulation layer between the electrode and dura^31, 12^. All models included dorsal root (DR) fibers as a cable model.

In 2002, Rattay and colleagues developed T11-L2 SCS model with basic precision in vertebrae, epidural space, dura, CSF, and thorax, whereas the spinal cord (including gray-matter and white-matter) had moderate precision in anatomical details. They modeled DR fibers as an electrical cable model^39^. Holsheimer in 2002 used a simplified 3D SCS model that lacked a dura mater compartment (basic precision) to discuss which nerve fibers (cable model of DR fibers) along the spinal cord were activated by SCS intensities within the therapeutic range ^10^. Manola and colleagues in 2005 and 2007 used a basic precision 3D SCS model that included vertebrae, epidural fat, dura, CSF, gray-matter, white-matter, and general thorax in the FEM. They also modelled DR fibers as a cable model ^40,41^. A more sophisticated computer-aided design (CAD)-derived SCS model was developed in 2010 by Ladenbauer and colleagues and Danner and colleagues in 2011 for non-invasive and invasive SCS with a vertebral column of moderate precision ^26,42^. This model represented spinal tissue compartments at basic anatomical precision, with uniform dimensions across spine levels. The model also did not include a dura mater and only had a single anterior root (AR) and posterior root (PR) fiber. These AR and PR fiber activation were further analyzed using a cable model ^26^. In 2011, Howard and colleagues developed a simplified 2D SCS model with no dura mater and limited precision in the modelled tissue compartments ^22^. Lee and colleagues in 2011 and Veizi and colleagues in 2017 developed a 3D FEM SCS model of a low thoracic and a sacral level spinal cord where most tissue compartments had basic anatomical precision, except the spinal cord (white-matter and gray-matter) which had an enhanced precision. They also modeled the electrical cable model of DR fibers ^13,30^. Hernández-Labrado and colleagues in 2011 developed a simplified C2-T1 SCS model comprising major spinal canal tissue compartments with basic anatomical precision ^21^. Parazzini and colleagues in 2014 and Fiocchi and colleagues in 2016 developed an MRI-derived SCS model for non-invasive SCS with enhanced anatomical precision in vertebrae, CSF, and spinal cord; however, other major spinal canal tissue compartments, such as epidural space, dura mater, and roots were not modelled. Moreover, it was unclear whether epidural fat was included in the FEM model ^20^. In 2013, Laird and colleagues developed a limited precision SCS model while modeling spinal root fiber as an electrical cable model ^43^. Howell and colleagues in 2014 constructed a SCS model (lower thoracic/upper lumbar spine level) with moderate anatomical precision in vertebrae, intravertebral disc, dural sac, CSF, white-matter, and gray-matter, and basic anatomical precision in soft tissue (thorax), with tissue compartment dimensions uniform across spine level ^23^. In 2014, Huang and colleagues developed a simplified 3D SCS model comprising major spinal canal tissue compartments with basic anatomical precision for epidural and intradural stimulation ^24^. In the same year, Kent and colleagues developed another simplified SCS model comprising vertebrae, epidural fat, dura mater, CSF, white-matter, and gray-matter, all with basic anatomical precision ^25^.

Lempka et al, 2015 developed a 3D SCS model (lower thoracic spinal cord) of kilohertz frequency spinal cord stimulation with white-matter and gray-matter with enhanced precision and other tissue compartments with basic anatomical precision ^44^. Miranda et al., 2016 developed the first MRI-derived SCS model comprised of nine tissue compartments; namely skin, fat plus subcutaneous adipose tissue, muscle, bone, dura mater (brain), vertebrae, IV discs, CSF, white-matter, and gray-matter. The model had enhanced anatomical precision on surrounding tissue compartments (vertebrae, IV Disc, and CSF), but moderate precision in dura, spinal cord (white-matter and gray-matter), and peripheral spinal tissues/thorax. It was unclear whether epidural space (fat) was included in the model ^27^. Arle and colleagues in 2014 and 2016 used an FEM derived from the Wesselink group’s ^37^ and Holsheimer group’s SCS model ^28,36^ (basic anatomical precision) ^16,45^. In their 2014 SCS model, vertebrae and spinal roots were missing ^45^.

Durá et al., 2019 developed a simplified 3D SCS model at the T10 spine level with basic anatomical precision in vertebrae, epidural fat, dura mater, CSF, while both white-matter and gray-matter had moderate anatomical precision. Modeled DR fibers had limited anatomical precision ^19^. In 2018, Kent and colleagues developed a 3D dorsal root ganglion (DRG) model with basic anatomical precision of the dorsal root and the DRG, and limited precision in epidural tissues and vertebrae ^46^. Wagner and colleagues in 2018 constructed a moderate precision SCS model of L1-S2 spine level including epidural fat, CSF, gray-matter, white-mater, spinal roots (dorsal and ventral), and rootlets. However, it was unclear whether vertebrae, discs, dura, and thorax were included in the model ^47^. Lempka and colleagues in 2019 constructed a patient-specific FEM SCS model with spinal cord, CSF, epidural fat, and a simplified spine domain. All segmented tissue compartments had moderate anatomical precision. They also modeled DR fibers’ as electrical model ^48,49^.

Anderson and colleagues in 2019 developed a simple 3D SCS model with basic anatomical precision of major spinal canal tissue compartments (white-matter, gray-matter, CSF, dura, and extra dural tissue layer) ^5^. Human L5 DRG model with basic anatomical precision was developed by Graham et al. in 2019 where they represented general thorax, bone, intraforaminal tissue, dural covering, and DRG using simplified shapes ^50^. Bikson’s group in 2019 developed a simplified T8-T10 SCS model (Generation 1 SCS model) comprising vertebrae (moderate precision), intra vertebral disc (IV Disc; moderate precision), epidural space/fat (basic precision), meninges/dura mater (basic precision), CSF (basic precision), spinal cord (basic precision), spinal roots (basic precision), rootlets (basic precision), thoracic aorta and sub-vasculature (basic precision), and soft tissues/thorax (basic precision) ^2^. In the same year, Lempka’s group developed an updated 3D SCS model of the lower thoracic spine level consisting gray-matter and white-matter of the spinal cord (enhanced anatomical precision), dorsal rootlets (moderate precision), CSF (basic precision), dura mater (basic precision), epidural fat (basic precision), vertebrae (moderate precision), and discs (moderate precision). This model had additional details and precision in some tissue compartments compared to their 2015 SCS model, but some major spinal tissue compartments were not included in the model, the dimensions of the tissue compartments were uniform across the spine level, and the surrounding tissue/thorax had simplified geometry ^15^.

These past studies clearly demonstrate that computational models represent a valuable tool to study the potential mechanisms of action of SCS and to optimize the design and implementation of SCS technologies. However, it is imperative that these computational models include the appropriate level of details to accurately predict the neural response to SCS and to correlate model predictions with clinical outcomes. Various simplifications to the model design may affect model-based predictions of the neural response to SCS. Therefore, we believe that there is a need for an anatomically-detailed high-resolution spinal cord model which captures major spinal tissue compartments in order to support enhanced prediction of SCS current flow.

## Methods

### State-of-the-art RADO SCS model

Adding more details and increasing resolution and anatomical precision, we developed the first Realistic Anatomically Detailed Open-source SCS (RADO-SCS) model. In this model, we included additional spinal tissue compartments that were not previously developed/modelled. Adding these extra tissues will influence the current flow pattern from the SCS lead to the spinal cord or to another possible region of interest. The dimensions and boundaries of tissue compartments of the RADO-SCS model were driven from human cadaver studies from lower thoracic spinal cord as discussed in our prior studies ^2,8,44,51^. In addition, some features such as Lissauer’s tract, thoracic aorta, sympathetic chains, dorsal and ventral roots, and rootlets were constructed based on physiological trajectory data ^52^. The RADO-SCS model consists of major spinal canal and peripheral tissue compartments with basic, moderate, and enhanced anatomical precision: vertebrae (moderate precision), intravertebral disc (moderate precision), epidural space (moderate precision), epidural space vasculature (basic precision), dura mater (moderate precision), dural sac (basic precision), intraforaminal tissue (basic precision), CSF (moderate precision), white-matter (enhanced precision), spinal cord vasculature (basic precision), Lissauer’s tract (enhanced precision), gray-matter (enhanced precision), dorsal and ventral root and rootlets (moderate precision), DRG (moderate precision), sympathetic chain (trunk and ganglion) (basic precision), thoracic aorta and its branching (basic precision), peripheral vasculatures (basic precision), and soft tissues (basic precision) (**Fig. 1**).

**Figure 1:**
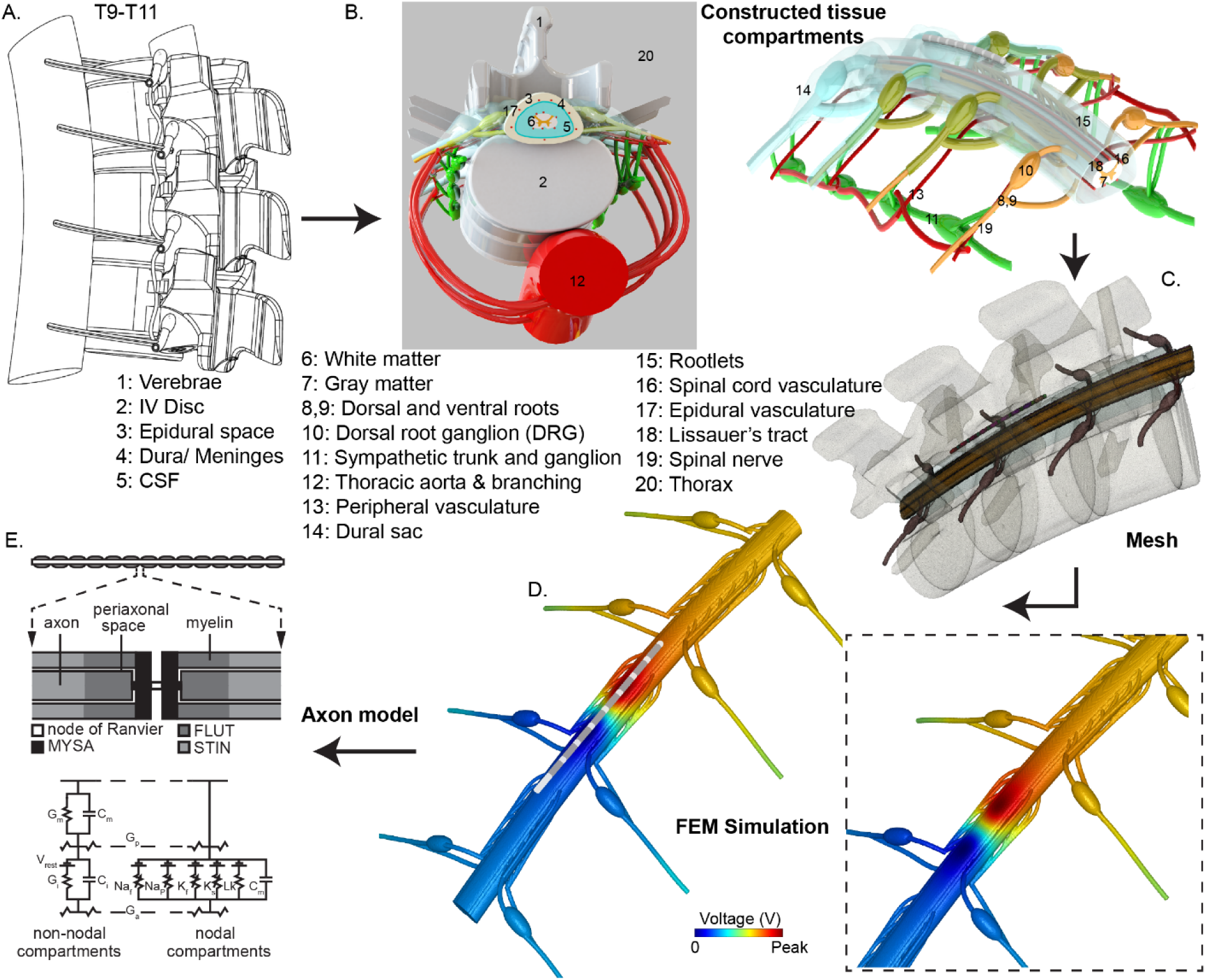
Computational FEM modelling and multi-compartment axon model pipeline of the RADO SCS. (A) An outline of the T9-T11 spinal cord CAD geometry. (B) Constructed tissue compartments of the detailed SCS model. (C) Final mesh with adequate mesh quality (D) FEM prediction of the spinal current flow (electric field). (E) Multi-compartment sensory axon model used to predict activation threshold for different fiber diameters using the voltage distribution at the surface of the spinal cord (panel modified from ^53^).

Specifically, we modelled and positioned 3 vertebrae and IV discs to mimic the T9-T11 lower thoracic spine with an anatomical curvature and tissue specific flexion. Four DRG were modelled lateral to the vertebrae (each side) in the rostro-caudal direction. The dorsal and ventral root converged together just beyond the DRG, while moving away from the cord to form a spinal nerve within the intervertebral foramen. These nerve and roots were surrounded by meninges and CSF.

The dorsal and ventral rootlets emerged from the dorsal and ventral horn of the spinal cord. We constructed eight dorsal and eight ventral rootlets at each spinal level. The thoracic aorta (which supplies arterial blood to the spinal cord) and its anastomotic network of radicular arteries that run along the posterior and anterior roots of the spinal nerves was constructed. The radicular arteries further branched at the spinal cord. Two sympathetic chains (trunk and ganglion) were constructed and the nerve from each sympathetic trunk was connected to the spinal nerve. Dural sac/covering, a membranous sheath that is part of the subarachnoid space, contains CSF, and surrounds the spinal cord, was also constructed. Just outside the dural sac was an epidural space, a major spinal tissue compartment which lies between dura mater and the vertebral wall, and predominantly contains fat. We modelled miniature blood vessels within the epidural space. Next, we constructed dura which is the outermost layer of the meninges. Inside the dura was a layer of conductive CSF which mimics the subarachnoid space that exists between the arachnoid and the pia mater. The inner most constructed tissue compartments were white-matter and gray-matter domains representing the spinal cord. We also constructed Lissauer’s tract, which was wedged between the dorsal horn and the surface of the spinal cord. Next, we placed the T9-T11 thoracic spinal column inside a thorax/soft tissues. We modeled an eight-contact clinical SCS lead (diameter: 1.25 mm, electrode contact length: 3 mm, inter-electrode insulation gap: 1 mm) (**Fig. 2**) and placed it 1 mm distal to the mediolateral dorsal column midline at the T10 spinal level.

**Figure 2:**
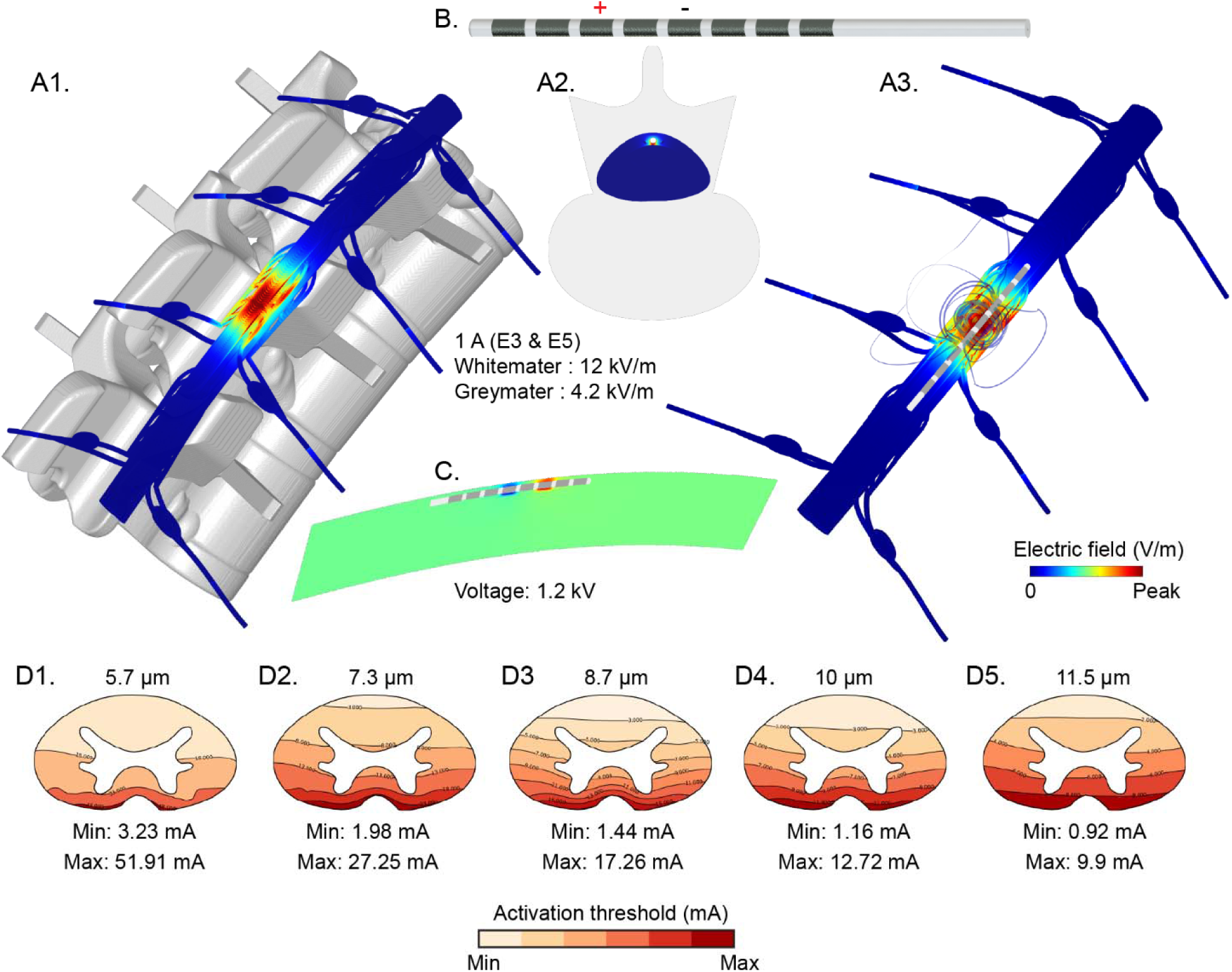
Predicted voltage distribution and electric field from the FEM and fiber activation thresholds. (A1, A2, A3) Predicted electric field at 1 A stimulation was 12 kV/m (white matter) and 4.2 kV/m (gray matter), respectively. (B) Illustration of the SCS lead used in the model. (C) Predicted peak voltage (1.2 kV) at the surface of the spinal cord. (D1, D2, D3, D4, D5) activation threshold for different fiber diameters. For 5.7, 7.3, 8.7,10.0, and 11.5 µm fiber diameters, the maximum activation thresholds were 51.9 mA, 21.3 mA, 17.3 mA, 12.7 mA, and 9.9 mA, respectively.

### Computational FEM model solution method

The assembled SolidWorks CAD model files along with the SCS leads (SolidWorks, Dassault Systemes Corp., MA, USA) were imported into Simpleware (Synopsys Inc., CA) to correct for some tissue specific anatomical anomalies (for e.g., overlapping, extrusion, smoothing) using morphological image processing filters. A volumetric FEM model was then generated from the final tissue masks. Using voxel-based meshing algorithms, an overly dense adaptive tetrahedral mesh of the SCS model was generated. After multiple mesh refinements that yielded within 1% error in voltage and current density at the spinal cord, a final mesh quality of greater than 0.9 was obtained (COMSOL mesh quality) indicating optimal elements (> 50 million tetrahedra elements). The volumetric mesh was later imported into COMSOL Multiphysics 5.1 (COMSOL Inc., MA, USA) to generate a FEM. We then solved the Laplace equation for electric current physics *(∇(σ∇V) = 0*) (V is potential and σ is electrical conductivity) under a steady-state assumption to determine the voltage distributions throughout the model. Assigned tissue and electrode conductivities were based on prior literature ^2,3,8,54^. Boundary conditions were applied in a bipolar configuration, with a 1 A load condition applied at electrode contact 3 (E3) while grounding electrode contact 5 (E5). Insulation (J.n = 0) on all remaining external boundaries of the model and continuity for the internal boundaries were assigned as other boundary conditions. We also assigned floating boundary conditions to the remaining inactive electrodes in the model. The relative tolerance was set to 1 × 10^−6^ to improve the solution accuracy. The 3D extracellular voltage distributions calculated from the FEM were exported and applied to the axon models described below.

### Multicompartment cable model of sensory Axons

Computer models of sensory axons within the dorsal columns of the spinal cord were developed based on previously published model of a mammalian sensory axon for specific fiber diameters. These fibers were parametrized to reproduce action potential shape, conduction velocity, and strength-duration relationship for sensory axons ^50^. Each sensory Aβ axon model was a double-cable model consisting of nodes of Ranvier separated by three distinct myelin segments: the myelin attachment segment (MYSA), paranodal main segment (FLUT) and the internode regions (STIN).

We distributed the sensory axon models throughout the white matter of the spinal cord using Lloyd’s Algorithm ^55^. The specific fiber sizes considered in our model matched the diameters explicitly parameterized in a previous study ^53^. We calculated the density of fibers in the model using histological measurements of fibers in the most superficial 300 µm of the dorsal columns^56^. To reduce computational demand, the total number of fibers solved for this project was reduced to 1% of anatomic density.

To determine the activation thresholds for each individual fiber, we applied the extracellular voltages calculated in the FEM to our axon models. We modeled the time-dependent output generated by an implantable pulse generated during current-controlled stimulation. To calculate the appropriate spatiotemporal voltage distributions, we then scaled the time-dependent voltage output by the spatial FEM voltage solution ^49,57^. We then interpolated the scaled extracellular voltages onto the model axons and used a bisection algorithm (error < 1%) to calculate the activation threshold for each axon. In our simulations, we applied a stimulus train consisting of pulses applied at a rate of 50 Hz, pulse width of 300 us, and a passive discharge phase of 6 ms in duration. We included a total of 3 pulses in our simulations. For each axon, we defined the activation threshold as the lowest pulse amplitude required to generate one action potential for each of the final 2 pulses of our 3-pulse stimulus train.

## Results

We used the RADO-SCS model to predict the voltage distribution and electric field in different tissue compartments with a clinical SCS lead positioned epidurally over the targeted lower thoracic vertebral column. Predicted peak electric field in the white matter and gray matter were 12 kV/m and 4.2 kV/m for 1A stimulation current, respectively. Peak predicted voltage at the surface of the spinal cord (white matter) was 1.2 kV (Fig. 2). The predicted electric field intensities were not uniform in the spinal tissues.

The activation thresholds throughout the white matter are shown in Fig. 2. As expected, the largest fibers had the lowest activation thresholds, and the most dorsal fibers were activated at thresholds below 1 mA. As the diameter of the fibers decreased, the activation thresholds increased; the smallest fibers in the model had a minimum activation threshold of 3.23 mA.

## Discussion

The SCS volume conductor models are systematically used to optimize clinical implementation of SCS technologies with ongoing efforts to enhance model precision and accuracy. Here, our goal is to develop an open-source high-resolution and anatomically-detailed SCS model and disseminate it to the scientific community. Currently, there is limited access to an open-source platform for SCS modeling, and currently-available have simple geometries and do not incorporate important tissue compartments. Thus, there is a need to develop and disseminate a high-resolution open-source SCS model. Using our modelling proficiencies, high-end computer resources, and an extensive literature search on anatomical details of spinal cord and peripheral tissues, we developed the first high-resolution open-source SCS model. This model is not only sophisticated in terms of details and architecture, but also has good precision to predict meaningful current flow. During the dissemination process, all the solid work part files and the assembly files, STL files, and volumetric mesh with/without a clinical SCS lead will be disseminated. In addition, any direct question regarding the files download or use will be answered by the corresponding authors via email. Any new updates will be periodically added to the source webpage.

## Acknowledgement

Source(s) of financial support: This study was partially funded by grants to MB from NIH (NIH-NINDS 1R01NS101362, NIH-NIMH 1R01MH111896, NIH-NCI U54CA137788/U54CA132378, and NIH-NIMH 1R01MH109289), PSC CUNY, Cycle 50.

